# Evolution of resistance to fluoroquinolones by dengue virus serotype 4 provides insight into mechanism of action and consequences for viral fitness

**DOI:** 10.1101/2020.05.11.088690

**Authors:** Stacey L. P. Scroggs, Jordan T. Gass, Ramesh Chinnasamy, Steven G. Widen, Sasha R. Azar, Shannan L. Rossi, Jeffrey B. Arterburn, Nikos Vasilakis, Kathryn A. Hanley

**Affiliations:** Department of Biology, New Mexico State University, Las Cruces, NM, USA; Department of Chemistry and Biochemistry, New Mexico State University, Las Cruces, NM, USA; Department of Biochemistry & Molecular Biology, The University of Texas Medical Branch, Galveston, TX, USA; Department of Pathology, The University of University of Texas Medical Branch, Galveston, TX, USA; Center for Biodefense and Emerging Infectious Diseases, The University of University of Texas Medical Branch, Galveston, TX, USA; Center for Tropical Diseases, The University of University of Texas Medical Branch, Galveston, TX, USA; Institute for Human Infection and Immunity, The University of University of Texas Medical Branch, Galveston, TX, USA

**Keywords:** Dengue virus, antiviral, fluoroquinolone, ciprofloxacin, enoxacin, difloxacin, evolution, resistance, fitness, mechanism-of-action

## Abstract

Drugs against flaviviruses such as dengue (DENV) and Zika (ZIKV) virus are urgently needed. We previously demonstrated that three fluoroquinolones, ciprofloxacin, enoxacin, and difloxacin, suppress replication of six flaviviruses. To investigate the barrier to resistance and mechanism(s) of action of these drugs, DENV-4 was passaged in triplicate in HEK-293 cells in the presence or absence of each drug. Resistance to ciprofloxacin was detected by the seventh passage and to difloxacin by the tenth, whereas resistance to enoxacin did not occur within ten passages. Two putative resistance-conferring mutations were detected in the envelope gene of ciprofloxacin and difloxacin-resistant DENV-4. In the absence of ciprofloxacin, ciprofloxacin-resistant viruses sustained a significantly higher viral titer than control viruses in HEK-293 and HuH-7 cells and resistant viruses were more stable than control viruses at 37°C. These results suggest that the mechanism of action of ciprofloxacin and difloxacin involves interference with virus binding or entry.

## INTRODUCTION

Members of the genus *Flavivirus*, most notably DENV and ZIKV, pose a great and growing threat to public health, but at present no approved antiviral therapies are available to treat any flaviviral infection (1–3). Drug repurposing offers the fastest route to move anti-flaviviral drugs into the clinic (4), and to this end we have recently demonstrated that three FDA-approved fluoroquinolones, enoxacin, difloxacin, and ciprofloxacin, suppress replication of six flaviviruses, including DENV and ZIKV, in cultured HEK-293 (human embryonic kidney) cells (5). Additionally, we found that treatment with enoxacin suppressed replication of ZIKV in the testes of interferon-deficient A129 mice *in vivo*, although it did not impact ZIKV titer in the serum, brain, or liver in these mice (5). Further, Xu et al. (6) recently reported that enoxacin also suppresses ZIKV replication in human neural progenitor cells (hNPCs) and brain organoids (6). Thus, fluoroquinolones offer a promising candidate for a repurposed therapy for flavivirus infections. However, several key aspects of the fluoroquinolone-flavivirus interaction must be elucidated before these drugs can move forward on the path to the clinic, particularly: (i) the barrier to resistance to these drugs, (ii) patterns of cross-resistance among fluoroquinolones, (iii) the fitness consequences of evolution of drug resistance, (iv) the mechanism(s)-of-action of fluoroquinolones against flaviviruses.

Resistance evolution has been a serious obstacle to successful treatment of other RNA viruses, particularly HIV, HCV, and influenza (7–9). Moreover, cross-resistance to structurally related drugs can occur and must be investigated as cross-resistance can render an entire class of antivirals ineffective (10,11). The utility of an antiviral also depends on the barriers to resistance (11–14). A drug that imposes a high genetic barrier for the evolution of resistance is more useful than a drug that allows for evolution of resistance to occur within a few passages or within a single host infection (13,15). RNA viruses have been shown to evolve drug resistance via one of two mechanisms: (i) specific antagonism of drug action, which often results in a decrease in fitness in the absence of the drug (9,16–21) or (ii) a general increase in fitness, which can sometimes be sustained following withdrawal of the drug (21,22). A decrease in fitness of resistant viruses could negatively impact viral transmission and alleviate disease burden, while a gain in fitness could exacerbate disease and could increase transmission, although these two phenotypes may also be uncoupled by evolution of drug resistance.

Previous studies have revealed multiple impacts of fluoroquinolones on both cellular and viral processes that may confer antiviral activity and shape the genetic pathway to resistance. First, multiple fluoroquinolones enhance RNA interference (23–25). Xu et al. (6) recently reported that enoxacin suppresses ZIKV in human neural progenitor cells (hNPC) and suggested the efficacy was due to enhancement of RNAi. Second, multiple fluoroquinolones have been shown to suppress hepatitis C virus (HCV), possibly by interfering with the viral helicase (26,27). Third, ciprofloxacin inhibits cellular helicases (28), which are targeted and bound by the flavivirus capsid protein (29) and the untranslated regions (30,31) to regulate viral replication and aid in virion assembly. Fourth, ciprofloxacin and trovafloxacin prevent cellular apoptosis (32,33), a process that flaviviruses suppress early in infection by activating the phosphatidylinositol 3-kinase pathway (34), but activate later in infection with the capsid, NS2A, and NS3 protease proteins (35–37). Fifth, ciprofloxacin suppresses autophagy (32). The loss of autophagy results in non-infectious virions and a decrease in viral RNA production during DENV infection (38). Lastly, the fluoroquinolone levofloxacin inhibits rhinovirus replication by reducing expression of the cellular receptor required for viral entry (39). Any one of these mechanisms, or combinations thereof, may be responsible for the efficacy of fluoroquinolones to suppress flaviviruses. There may also be functions of fluoroquinolones that have not been described in the literature that contribute to their anti-flaviviral efficacy. The location of mutations specific to fluoroquinolone-resistant viruses can suggest possible mechanisms-of-actions.

In the current study, we harnessed the power of experimental evolution to investigate fluoroquinolone-flavivirus interactions. We hypothesized that: (i) the barrier to resistance against fluoroquinolones would be low and that, therefore, resistance would evolve within ten passages in cultured cells, (ii) resistance to one fluoroquinolone would confer cross-resistance to other fluoroquinolones due to the high structural similarity of these drugs, and (iii) resistance would evolve via specific evasion of the drug, resulting in a decrease in fitness in the absence of the drug. To test these hypotheses, we serially passaged DENV-4 in triplicate in the presence or absence of fluoroquinolones, in human embryonic kidney cells (HEK-293). While DENV-4 evolved resistance to ciprofloxacin in 7 passages and difloxacin in 10 passages, it did not evolve resistance to enoxacin in 10 passages. We predicted that ciprofloxacin-resistant and control lineages of DENV-4 would share common mutations relative to the parent DENV-4 that confer adaptation to HEK-293 cells (40), but that ciprofloxacin-resistant lineages would also possess a unique constellation of mutations, some conferring fluoroquinolone-resistance, and that these latter mutations would provide insight into the fluoroquinolone mechanism-of-action (41–50). We also used the ciprofloxacin and difloxacin-resistant DENV-4 lineages to test the hypothesis that evolution of resistance to one fluoroquinolone will confer cross-resistance to other members of this class. Finally, we quantified the fitness and stability of resistant, control and the parent virus in the absence of drug in mammalian and mosquito cultured cells and mosquitoes *in vivo* to draw inference about the mechanisms underlying resistance. Evolutionary studies suggest that gains in flavivirus fitness in the mammalian host come at the cost of a loss in fitness in the mosquito vector (a trade-off) (51–53). As both fluoroquinolone-resistant and media-control DENV-4 had been passaged in HEK-293 cells, we predicted that these lineages would both have lower fitness in mosquitoes than the parent DENV-4, and that the ciprofloxacin-resistant virus would be more disadvantaged than the media-control virus. In sum, these studies seek to move the evaluation of FDA-approved drugs for antiviral repurposing beyond the demonstration of efficacy to consider the consequences of resistance.

## MATERIALS AND METHODS

### Viruses and cells

Dengue virus serotype 4 (rDEN-4 Dominica p4) was derived from full-length clone p4 (54) and passaged four times in Vero cells. Replicate vials of a working stock were collected in 1X SPG (2.18 mM sucrose, 38 mM potassium phosphate [monobasic], 72 mM potassium phosphate [dibasic], 60 mM L-glutamic acid), clarified by centrifugation, and stored at −80 °C. HEK-293 (human embryonic kidney) cells were obtained from ATCC (CRL-1573); Vero, HuH-7, and C6/36 cells were obtained from the lab of Dr. Stephen Whitehead (NIAID, NIH). HEK-293 cells were maintained at 37°C with 5% CO_2_ in Dulbecco’s minimum essential medium (DMEM/F12; Gibco, Life Technologies, Grand Island, NY) supplemented with 10% heat-inactivated fetal bovine serum (FBS) (Gibco), 2mM L-glutamine (Gibco), and 0.5% antibiotic-antimycotic (penicillin, streptomycin, and amphotericin B; Gibco). Vero cells were maintained at 37°C with 5% CO_2_ in minimum essential medium (MEM, Gibco) supplemented with 10% heat-inactivated FBS, 2mM L-glutamine, and 0.05 mg/mL gentamycin (Gibco). HuH-7 cells were maintained at 37°C with 5% CO_2_ in DMEM/F12 supplemented with 10% heat-inactivated FBS, 2mM L-glutamine, and 0.05 mg/mL gentamycin. C6/36 cells were maintained at 32°C with 5% CO_2_ in MEM supplemented with 10% heat-inactivated FBS, 2mM L-glutamine, 2mM nonessential amino acids (Gibco), and 0.05 mg/mL gentamycin. Viral titers were determined via serial dilution onto HEK-293, Vero, HuH-7 or C6/36 cells followed by immunostaining using previously described methods (5,55,56). Briefly, each virus was serially diluted ten-fold and inoculated onto confluent cells in 24-well plates. After two hours of incubation at 37°C with occasional rocking, infected cells were overlaid with 1% methylcellulose in OptiMEM (Gibco) that had been supplemented with 2% FBS, 2mM L-glutamine, and 0.05 mg/mL gentamycin. Plates were incubated for five days under maintenance conditions, after which cells were fixed with ice cold methanol: acetone (1:1) for HEK-293 and 90% methanol for Vero, HuH-7, and C6/36 cells for 30 minutes. Viral plaques were immunostained using DENV-4-specific hyperimmune mouse ascitic fluid and peroxidase-labeled goat anti-mouse secondary antibody (KPL, Gaithersburg, MD) then developed with KPL True Blue Peroxidase Substrate (SeraCare, Milford, MA) and counted to calculate viral titer.

### Antiviral compounds

For each experiment, a fresh working stock of enoxacin (Sigma-Aldrich, E3764, St. Louis, MO), difloxacin (Sigma-Aldrich, D2819, St. Louis, MO), or ciprofloxacin (Corning, 86393-32-0, Manassas, VA) at a concentration of 1.5 mM was sonicated in nanopore water with 3 mM lactic acid (Sigma-Aldrich, L1750, St. Louis, MO) and sterilized *via* passage through a 0.2 μm filter as described in (5). The compounds were diluted to their final concentrations in HEK-293 cell culture media.

### Selection of drug-resistant viruses

Fluoroquinolone-resistant DENV-4 was generated by passaging the virus in triplicate in the presence of enoxacin, difloxacin, or ciprofloxacin (three independent lineages per drug) until resistance was detected (Figure 1a, Figure 1c, Figure 1e) or for a total of ten passages. This endpoint was chosen based on selection for flavivirus resistance to other candidate antivirals in cell culture, in which a range of 7 to 21 passages was required to detect resistance (19,41,44–48,57), though we recognize that studies that did not detect resistance may not be in the published literature due to negative publication bias (58). Additionally, one set of triplicate control lineages were passaged in cell culture media. For each passage, T25 flasks of HEK-293 cells at 80% confluency were infected at a multiplicity of infection (MOI) of 0.1. After two hours incubation at 37°C with 5% CO_2_, infected cells were washed with 1x PBS and treated with 5 mL of each drug at the specified concentrations or media for each passage (Figure 1a, 1c, and 1e). The effective concentration 50 (EC_50_) values for difloxacin and ciprofloxacin, 10.1μM, and 19.6μM respectively as reported in (5) were used to initiate passage, however, the concentration of ciprofloxacin and difloxacin was increased 2-fold after 3 or 4 passages, respectively, to accelerate evolution of drug resistance. Based on preliminary experiments, passaging with enoxacin was initiated at 2x the EC_50_ or 15.2μM. When viral titers following drug treatment were not high enough to initiate the next passage (enoxacin passages 3 and 6 and difloxacin passage 8), the passage was repeated at half the previous drug concentration. All drug concentrations remained below the cytotoxic concentration 50 (CC_50_) of 1504.0μM, 763.0μM, 537.6μM for ciprofloxacin, difloxacin, and enoxacin in HEK-293 based on (5). After 5 days of incubation, viral supernatants were collected in 1x SPG, clarified by centrifugation, and stored at −80 °C. Viral titers were determined by plaque assay in HEK-293 cells.

**Figure 1.**
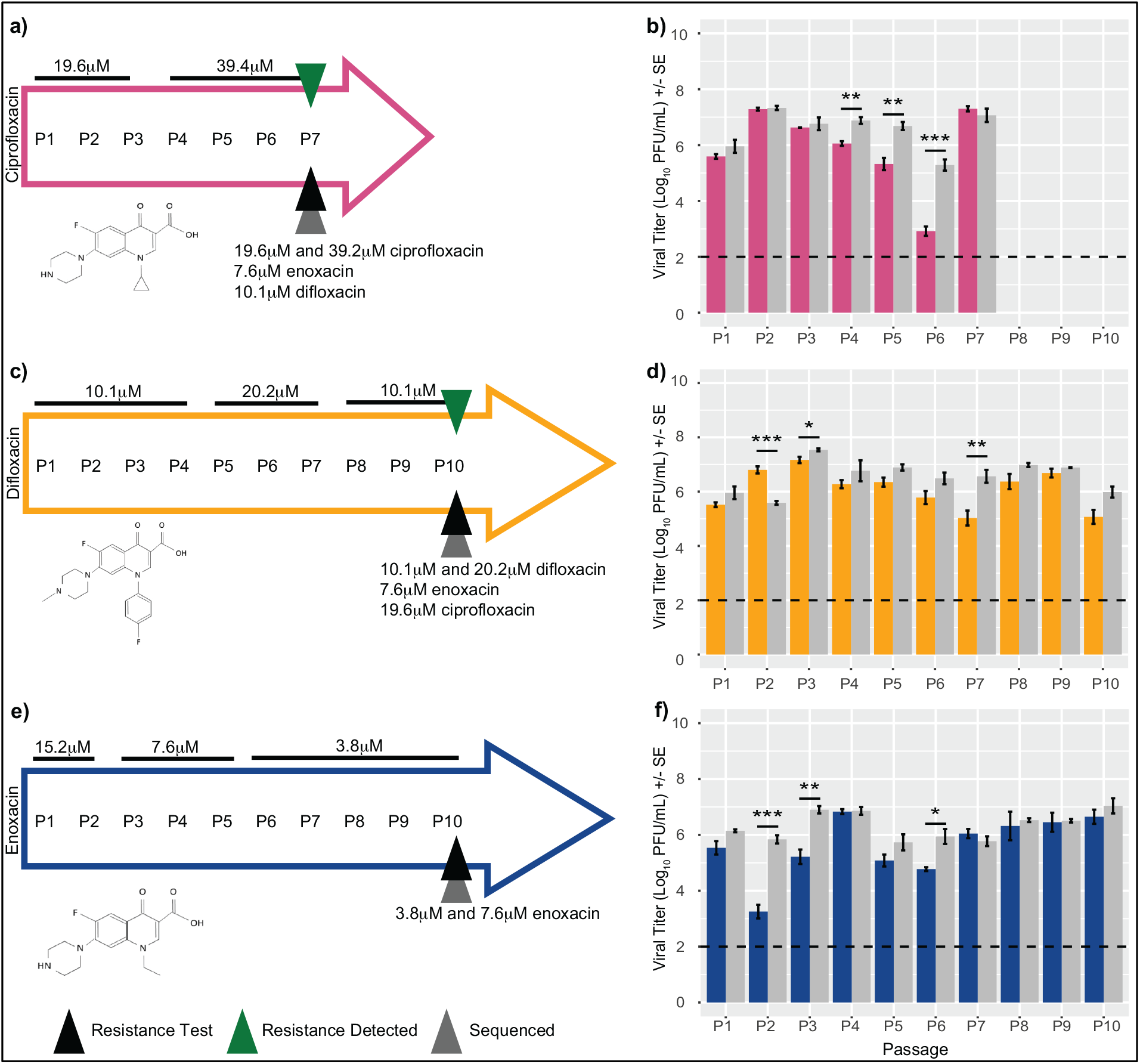
DENV-4 was passaged in the presence of ciprofloxacin (a), difloxacin (c), and enoxacin (e), at indicated concentrations for 10 total passages or until resistance was detected. Black triangles indicate the passage at which formal resistance tests were conducted at specified concentrations, green triangles indicate when resistance was confirmed, and grey triangles indicate when the genomes (for the ciprofloxacin treatment) or the structural genes (for the difloxacin and enoxacin treatments) were sequenced. Chemical structures of designated fluoroquinolones are depicted. Mean viral titers +/− SE by passage for DENV-4 passaged with media control (grey bars, b,d,f) or ciprofloxacin (b, pink bars), difloxacin (d, yellow bars), and enoxacin (f, blue bars). Dashed line indicates the limit of detection. * P < 0.05

### Antiviral resistance assays

Resistance was suspected when the mean viral titers of the fluoroquinolone-passaged viruses and the media-control viruses did not differ, or the titer of the fluoroquinolone-passaged viruses increased above the media-controls. When resistance to a specific fluoroquinolone was suspected, or after a total of ten passages, resistance was formally tested by infecting duplicate T25 flasks of HEK-293 cells with each of the three lines of virus passaged in that fluoroquinolone, the three control lines, or the parent virus (N = 7 virus lines) at MOI: 1.0. Two hours after infection, one flask of infected cells per virus was treated with the specific fluoroquinolone at the EC_50_ and twice the EC_50_ (ciprofloxacin: 19.6 μM and 39.2 μM, difloxacin: 10.1μM and 20.2 μM) or half the EC_50_ and the EC_50_ (enoxacin: 3.8 μM and 7.6 μM) in 5 mL media and the other flask was treated with 5 mL media. After five days, the viral supernatants were collected, clarified, frozen, and viral titers determined via plaque assay.

Ciprofloxacin-resistant viruses were tested for cross-resistance to enoxacin and difloxacin at 7.6μM and 10.1μM, respectively, and difloxacin-resistant viruses were tested for cross-resistance to enoxacin and ciprofloxacin at 7.6 μM and 19.6 μM, respectively. To test for cross-resistance, triplicate T25 flasks of HEK-293 cells were infected with each of the three lines of viruses resistant to the specified fluoroquinolone, the three control lines, or the parent virus (N = 7 virus lines) at MOI: 1.0. Two hours after infection, one flask of infected cells per virus was treated with the first target drug in 5 mL media, the next infected flask was treated with the second target drug in 5 mL media, and the last infected flask was treated with 5 mL media. After five days, the viral supernatants were collected, clarified, frozen, and viral titers determined via plaque assay.

### Library preparation and high-throughput sequencing of ciprofloxacin-resistant, media-control, and parent DENV-4

Whole genomes from the ciprofloxacin-resistant and media-control viruses from passage 7 were deep sequenced at the University of Texas Medical Branch at Galveston, Texas. Libraries for sequencing were prepared with the NEBNext Ultra II RNA Prep Kit (New England BioLabs, Inc.) following the manufacturer’s protocol. Briefly, ~70-100 ng of RNA was fragmented for 15 minutes, followed by cDNA synthesis, end repair and adapter ligation. After 8 rounds of PCR the libraries were analyzed on an Agilent Bioanalyzer and quantified by qPCR. Samples were pooled and sequenced with a paired-end 75 base protocol on an Illumina (Illumina, Inc) NextSeq 550 using the High-Output kit.

Reads were processed with Trimmomatic v0.36 (59) to remove low quality base calls and any adapter sequences. The *de novo* assembly program ABySS v1.3.7 (60) was used to assemble the reads into contigs, using several different sets of reads, and kmer values from 20 to 40. Contigs greater than 400 bases long were compared against the NCBI nucleotide collection using BLAST. A nearly full length DENV-4 viral contig was obtained in each sample. All the remaining contigs mapped to either host cell ribosomal RNA or mitochondria. The trimmed reads from each sample were mapped to the sample consensus sequence with BWA v0.7.17 (61) and visualized with the Integrated Genomics Viewer v2.3.26 (62) to confirm a correct assembly.

For single nucleotide variant and insertion/deletion calling the trimmed reads from each sample were mapped to the parental reference sequence (54) with BWA. The LoFreq v2.1.3.1 (63) call and call-indels commands were used for variant calling, after the mapped reads were preprocessed with the LoFreq viterbi and indelqual commands to fix alignments at the read ends and insert indel quality scores, respectively. Variant calls were filtered at a level of 0.5%. Shannon entropy was calculated for each replicate (64).

### RNA isolation, RT-PCR, and Sanger sequencing of difloxacin-resistant, enoxacin-passaged, and parent DENV-4

To probe the structural genes for the anti-flaviviral mechanism-of-action, the capsid, prMembrane, and envelope genes of the difloxacin-resistant, enoxacin-passaged, media-control DENV-4 passaged 10 times, and parent virus were sequenced (n = 10 viruses). Viral RNA was isolated using the QIAamp Viral RNA Mini Kit (Qiagen, Hilden, Germany) per the manufacturer’s protocol. RNA was reverse transcribed and amplified using previously validated (55) forward (17F: 5’ GGACCGACAAGGACAGTTCCAAAT 3’) and reverse (2546R: 5’ TCTCGCTGGGGACTCTGGTTGAAAT 3’) primers (Eurofins, Louisville, KY) and SuperScript™ III One-Step RT-PCR kit with Platinum™ Taq DNA Polymerase (Invitrogen, Carlsbad, CA) using the following thermocycler conditions: 50 °C for 30 min; 94 °C for 3 min; 9 cycles at 94 °C for 10 s, 60 °C for 30 s, and 68 °C for 4 m; 29 cycles at 94 °C for 10 s, 60 °C for 30 s, and 68 °C for 4 m with 5 s added at each cycle; and a final extension at 68 °C for 7 min. Amplicons were sequenced via Sanger sequencing at the Genomic Analysis Core Facility at the University of Texas at El Paso using the amplification primers listed above as well as four validated internal primers (996R: 5’ ATGCTCCACCTGAGACTCCTTCC 3’, 860F: 5’ GGCTTATATGATTGGGCAAACAGG 3’, 1683R: 5’ CCTGTCTCTTGGCATGAGGAACC 3’, 1551F: 5’ ACATGGCTCGTGCATAAGCAATGG 3’) (55). The resulting forward and reverse sequences were clipped for quality and aligned in Geneious (version 9.0.5 (65)) with the reference genome (GenBank AY648301.1). Mutations were identified in the passaged viruses compared to the parent DENV-4.

### Replication kinetics

To evaluate the impacts of ciprofloxacin resistance on DENV-4 replication dynamics in human and mosquito cells, 7 T25 flasks of 80% confluent HEK-293, HuH-7, Vero, or C6/36 cells were infected at MOI 0.1, based on the viral titer determined in HEK-293, with each of the ciprofloxacin-resistant DENV-4 or media control DENV-4 lineages (both from passage seven) or the parent DENV-4. After 2 hours of incubation with occasional rocking, the flasks were washed with 1X PBS and 5ml of cell-specific media was added to each flask. Viral supernatants were collected on the day of infection as well as 1, 2, 3, 4, 6, and 8 days post-infection (dpi). For replication curves with HuH-7 and Vero cells, the time points for collection were extended to include 9, 10, 11, and 12 dpi as the plateau viral titer was not detected by 8 dpi. At each time point, 1 ml of the media was removed from each flask and 1 ml of fresh media was added back to each flask. The viral supernatants were clarified by centrifugation at 1200 rpm for 10 minutes at 4°C, stored at −80 °C in 1X SPG, and viral titers were determined using the same cell line as that used for the replication dynamics (HEK-293, HuH-7, Vero, or C6/36).

### Stability assay

To detect differences in virion stability between the ciprofloxacin-resistant and media control DENV-4, 7 T25 flasks of 80% confluent HEK-293 cells were infected at MOI: 0.1 with each of the ciprofloxacin-resistant DENV-4 or media control DENV-4 lineages (both from passage seven) or the parent DENV-4. After 2 hours of incubation with occasional rocking, the flasks were washed with 1X PBS and 5ml of cell-specific media was added to each flask. On the expected day of peak viral titer. day 3, the viral supernatants were collected on ice then clarified by centrifugation at 1200 rpm for 10 minutes at 4°C. Immediately after centrifugation, 45 μL of each clarified supernatant was added to cryotubes with 405 μL Leibovitz L15 media (Gibco, Life Technologies, Grand Island, NY) and placed into a 37 °C incubator, one tube for each virus and for each time point (0 hours post-incubation, 2, 4, 6, 8, 12, 24, and 30). At each time point, the cryotube was removed from the incubator and 50 μL of 10X SPG was added before the tube was snap frozen on a dry ice-methanol bath. Frozen samples were stored at −80 °C and viral titers were determined using HEK-293 cells as described above.

### Mosquito Infection

Eggs from *Aedes aegypti* (Rockefeller strain) were hatched and reared as described in (66). Infectious bloodmeals were prepared and mosquito feeding was conducted as described in (56). Briefly, 1 mL of virus was mixed with 2 mL of washed red blood cells from defibrinated rabbit blood (Hemostat, Dixon, CA, USA) in 10% sucrose; 75 μl of 120 mM ATP was added to each meal immediately prior to feeding. Sealed cartons containing ~30 mosquitoes were starved for 24 hours, placed under parafilm-covered feeders warmed to 37°C, and allowed to feed for 20 minutes. Engorged mosquitoes were separated into new cartons and incubated at 27 °C with 80% relative humidity on a 12-hour light and 12-hour dark cycle. The mosquitoes were provided 10% sucrose in cotton pledgets *ad libitum*. After 10 days, mosquitoes were cold-killed and stored at −80 °C, after which whole mosquitoes were homogenized in Hank’s balanced salt solution (Gibco) supplemented with 10% FBS, 250μg/mL amphotericin B (Gibco), 1% ciprofloxacin, and 150 μg/mL clindamycin and viral titer was determined as described above in C6/36 cells with the addition of 5 μg/mL amphotericin B to the methylcellulose overlay media.

### Statistical analyses

Viral titer data were log transformed, assessed for normality and then compared using t-tests or two-way ANOVAs as appropriate. Repeated measures ANOVA was used to detect differences in replication kinetics and percent change in viral titer from the stability assay between the ciprofloxacin-resistant and media control viruses. The parent DENV-4 was not included in the repeated measures ANOVA as there was only one parental replicate. Differences in variant frequency, Shannon entropy, and mosquito mortality (%) were detected by ANOVA. Differences in variant frequency within genes and between treatments were identified using a two-way ANOVA. Differences in the counts of mosquito bodies positive for DENV-4 infection was detected using contingency table analyses. Tukey-Kramer or pairwise t-tests, as specified, were used to ascertain *post hoc* pairwise differences. Statistics were conducted in R using packages: tidyverse, plyr, dplyr, lme4, car, emmeans, lsmeans, and pbkrtest.

## RESULTS

### Evolution of resistance to each of three FQs by DENV

In this study, resistance was defined as a lack of a difference between the mean viral titers of the media-passaged control viruses cultured in media and viruses passaged in a specific fluoroquinolone cultured in the presence of that fluoroquinolone. After seven passages with increasing concentration of ciprofloxacin (Figure 1a, 1b), DENV-4 resistance to ciprofloxacin was detected (Figure 2a). An interaction between the passage condition (media or ciprofloxacin) and resistance test condition (media or ciprofloxacin) was found with a two-way ANOVA (F (1,8): 40.7, P = 0.0002). Treatment of both media-passaged virus and ciprofloxacin-passaged virus with 39.4μM ciprofloxacin (2x the EC_50_) resulted in a significant decrease in titer for both virus lineages relative to treatment with media (Tukey pairwise comparisons, P < 0.05), but the decrease in viral titer of the media-control viruses after ciprofloxacin treatment was greater than that of the ciprofloxacin-passaged viruses (4.2 log decrease vs 1.9 log decrease). There was no significant difference in mean viral titers between the media-passaged viruses treated with media and the ciprofloxacin-passaged viruses treated with ciprofloxacin (Tukey pairwise comparison, P < 0.05). In the absence of ciprofloxacin, replication of the ciprofloxacin-passaged viruses was not different than the media-passaged viruses (Tukey pairwise comparison, P > 0.05), and in the presence of ciprofloxacin, the ciprofloxacin-passaged viruses replicated to significantly higher levels (2.8 log increase) than the media-passaged viruses (Tukey pairwise comparison, P < 0.05).

**Figure 2.**
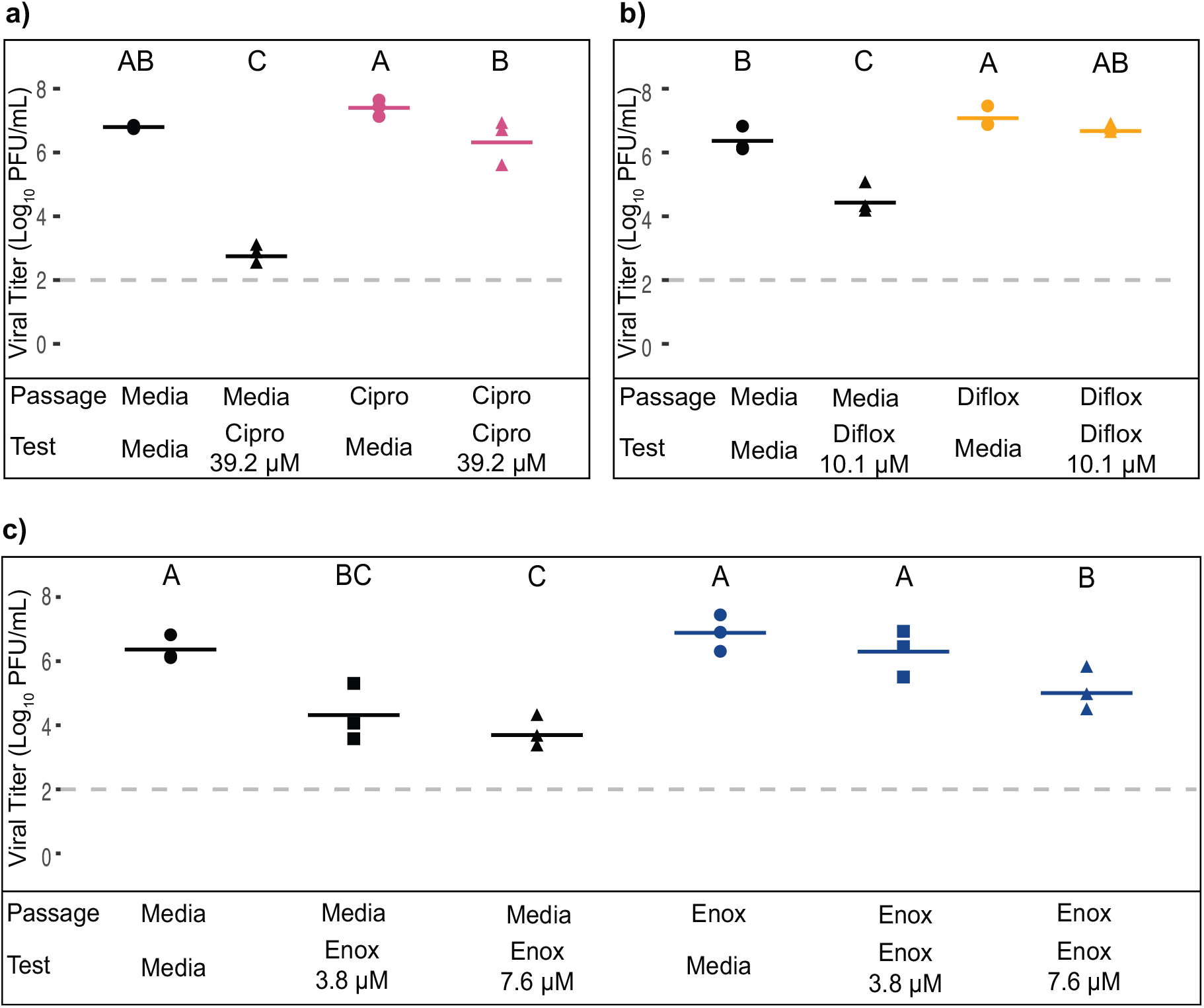
DENV-4 resistance to ciprofloxacin was detected after 7 passages (a), and difloxacin after 10 passages (b), but not to enoxacin (c). Black: media control passaged DENV-4, Pink: ciprofloxacin-passaged DENV-4, Yellow: difloxacin-passaged DENV-4, Blue: enoxacin-passaged DENV-4, Circles: Tested in resistance test with media, Triangles: Tested in resistance test with 39.2μM (x2 EC_50_) ciprofloxacin (a), 10.1 μM (EC_50_) difloxacin (b), 3.8 μM (1/2 EC_50_), 7.6 μM (EC_50_) enoxacin (c). Differences were detected with two-way ANOVA with Tukey pairwise comparisons (see text for full statistics). Solid lines indicate mean and dashed line indicates the limit of detection.

After ten passages (Figure 1c, 1d), DENV-4 had evolved resistance to a lower concentration (10.1 μM) of difloxacin (Figure 2b), but resistance was not detected when twice the concentration of difloxacin was tested (Figure S1). In the resistance test with the lower concentration of difloxacin, an interaction was detected between passage and resistance test conditions using a two-way ANOVA (F (1,8) = 13.7, P = 0.006) (Figure 2b) that was absent in the resistance test at the higher concentration of difloxacin (two-way ANOVA F (1,8) = 0.07, P = 0.8) (Figure S1). When the media-passaged viruses were treated with 10.1 μM difloxacin a significant reduction in viral titer occurred compared to the media treatment that was not detected when the difloxacin-passaged viruses were treated with difloxacin or media (Figure 2b; Tukey pairwise comparison, P < 0.05). Media and difloxacin-passaged viruses were both suppressed by 20.2μM (2x the EC_50_) difloxacin to at or below the level of detection. There was not a significant difference between the mean viral titers of the media-passaged viruses treated with media and the difloxacin-passaged viruses treated with difloxacin (Figure 2b; Tukey pairwise comparison, P > 0.05).

After DENV-4 was passaged 10 times in the presence of enoxacin (Figure 1e, 1f) the interaction between the passage condition (media or enoxacin) and resistance test condition (media or enoxacin at 3.8 μM or 7.6 μM) was not significant (two-way ANOVA F (1,2): 1.9, P = 0.19; Figure 2c). The mean viral titer for media-passaged viruses treated with media was not significantly different from the mean viral titer for the enoxacin-passaged viruses treated with 3.8 μM, or half the EC_50_, but was significantly different from the enoxacin-passaged viruses treated with the EC50 for enoxacin (7.6 μM) (Tukey pairwise comparison, P < 0.05; Figure 2c). These data indicate a trend toward enoxacin resistance but did not satisfy our *a priori* criteria for resistance.

### Mutations associated with fluoroquinolone resistance

Only two coding mutations became predominant (> 50% of all reads) in the ciprofloxacin-resistant but not in the media control viruses; these included a V15L mutation in domain I of the envelope glycoprotein (E) in two replicates of ciprofloxacin-resistant viruses and an E417A mutation in domain III of E in one replicate of the ciprofloxacin-resistant viruses (Table 1). One of these mutations, E417A, was also found in one replicates of difloxacin-resistant viruses (Table 2). Other coding mutations in the E, NS2B and NS4B genes and non-coding mutations in the untranslated regions (UTRs), many of which had been previously identified in the literature as HEK-adapting mutations, were found in at least a subset of both drug-resistant and control viruses (Table 1). No mutations were identified in the parent DENV-4 compared to the reference sequence (AY648301). Silent mutations within the open reading frame are listed in Table S1.

**Table 1.** Nucleotide and amino acid changes for coding mutations identified through deep sequencing in the ciprofloxacin-resistant, media-control DENV-4 from passage 7 as well as parent DENV-4

| Gene | Nucleotide change | Amino acid change | Present in ciprofloxacin-resistant DENV-4 (P7) |  |  | Present in media control (P7) |  |  | Parent DENV-4 | Possible function |
| --- | --- | --- | --- | --- | --- | --- | --- | --- | --- | --- |
|  |  |  | A | B | C | A | B | C |  |  |
| 5' UTR | A26G | NA | - | - | - | X | - | - | - | HEK293 adaptation |
|  | A31G | NA | X | - | - | - | - | - | - | HEK293 adaptation |
|  | U40C | NA | - | - | X | - | - | X | - | HEK293 adaptation |
| E | G981C | V15L | - | X | X | - | - | - | - | Ciprofloxacin resistance |
|  | A1918G | E327G | X | X | X | X | X | X | - | HEK293 adaptation (70,73) |
|  | A2188C | E417A | X | - | - | - | - | - | - | Ciprofloxacin resistance |
| NS2B | C4387U | S85F | X | X | - | - | X | - | - | HEK293 adaptation |
| NS4B | U6864C | F13L | X | X | - | - | X | - | - | HEK293 adaptation |
|  | C7129U | P101L | - | - | X | X | - | X | - | HEK293 adaptation (71) |
|  | C7141U | T105I | X | X | - | - | X | - | - | HEK293 adaptation (72) |
|  | G7182C | G119R | - | - | - | - | X | - | - | HEK293 adaptation |
|  | C7546U | A240V | - | X | X | - | - | X | - | HEK293 adaptation (40) |
| 3' UTR | U10290C | NA | X | X | - | - | X | - | - | HEK293 adaptation |
|  | U10352C | NA | X | X | - | - | X | - | - | HEK293 adaptation |

X: Mutation present at at least 0.5 frequency
-: Mutation not present at at least 0.5 frequency

**Table 2.** Nucleotide and amino acid changes for envelope gene identified with Sanger sequencing in the difloxacin-resistant, enoxacin passaged, media-control DENV-4 from passage 10 as well as parent DENV-4

| Gene | Nucleotide change | Amino acid change | Present in difloxacin-resistant DENV-4 (P10) |  |  | Present in enoxacin passaged DENV-4 (P10) |  |  | Present in media control (P10) |  |  | Parent DENV-4 | Possible function |
| --- | --- | --- | --- | --- | --- | --- | --- | --- | --- | --- | --- | --- | --- |
|  |  |  | A | B | C | A | B | C | A | B | C |  |  |
| E | A1918G | E327G | X | X | X | X | - | X | X | X | X | - | HEK293 adaptation (70,73) |
|  | A2188C | E417A | - | X | - | - | - | - | - | - | - | - | Difloxacin resistance |

X: Mutation detected
-: Mutation not detected

To assess amino acid conservation across the flaviviruses at positions 15 and 417 in E, we generated a multiple sequence alignment of mosquito-borne, tick-borne, no known vector, and insect-only flaviviruses (Figure S2). The valine as position 15 is conserved across most mosquito-transmitted flaviviruses, but a leucine occurs in this position in DENV-1 and DENV-3, which recreates the V15L mutation observed in the ciprofloxacin resistant DENV-4 in this study. Position 417 is a glutamic acid in DENV-4 and the tick-borne flaviviruses, an aspartic acid in DENV-1, DENV-2, DENV-3 and the other mosquito-transmitted viruses analyzed, a serine in the two tick-borne flaviviruses analyzed and a threonine in the single insect-only flavivirus analyzed. Notably, an alanine was not detected in this position in any of the viruses analyzed.

### Evolution to ciprofloxacin does not increase interpopulation viral diversity

Overall there was no consistent difference between the ciprofloxacin-resistant, media control or parent virus lineages in diversity. Mean frequency of variants was significantly different across the seven virus populations (Figure 3a; ANOVA: F(6, 394) = 3.51, P = 0.002); however pairwise comparisons revealed that the mean variant frequency of only one ciprofloxacin-resistant replicate (CB) was significantly higher than the parent population (6.0-fold increase) and one media control replicate population (MC) (3.3-fold increase) (Figure 3a). Genetic complexity, evaluated using Shannon entropy, was also different for the 7 populations (Figure 3b; ANOVA: F (6, 394) = 3.03, P = 0.007), but pairwise comparisons indicate that the only significant difference was between two ciprofloxacin-replicates (Figure 3b). The variants for all three viral populations were distributed across the genome (Figure 4). The interaction between gene location and viral population (ciprofloxacin-resistant, media control, or parent) in relation to mean variant frequency was not significant overall (repeated measures ANOVA: F(22, 365) = 1.12, P = 0.32), however pairwise analyses within genes show that the mean variant frequency of the ciprofloxacin-resistant populations was higher than that of the parent in the 5’ UTR (5.1-fold increase), envelope (12-fold increase), and NS4B (3.8-fold increase), but was only also significantly higher than the media control in NS4B (2.1-fold increase) (Figure 5, Table S2). The mean variant frequency for the ciprofloxacin-resistant populations was also higher than the media control population in NS2B but was not different than the parent (Figure 5, Table S2).

**Figure 3.**
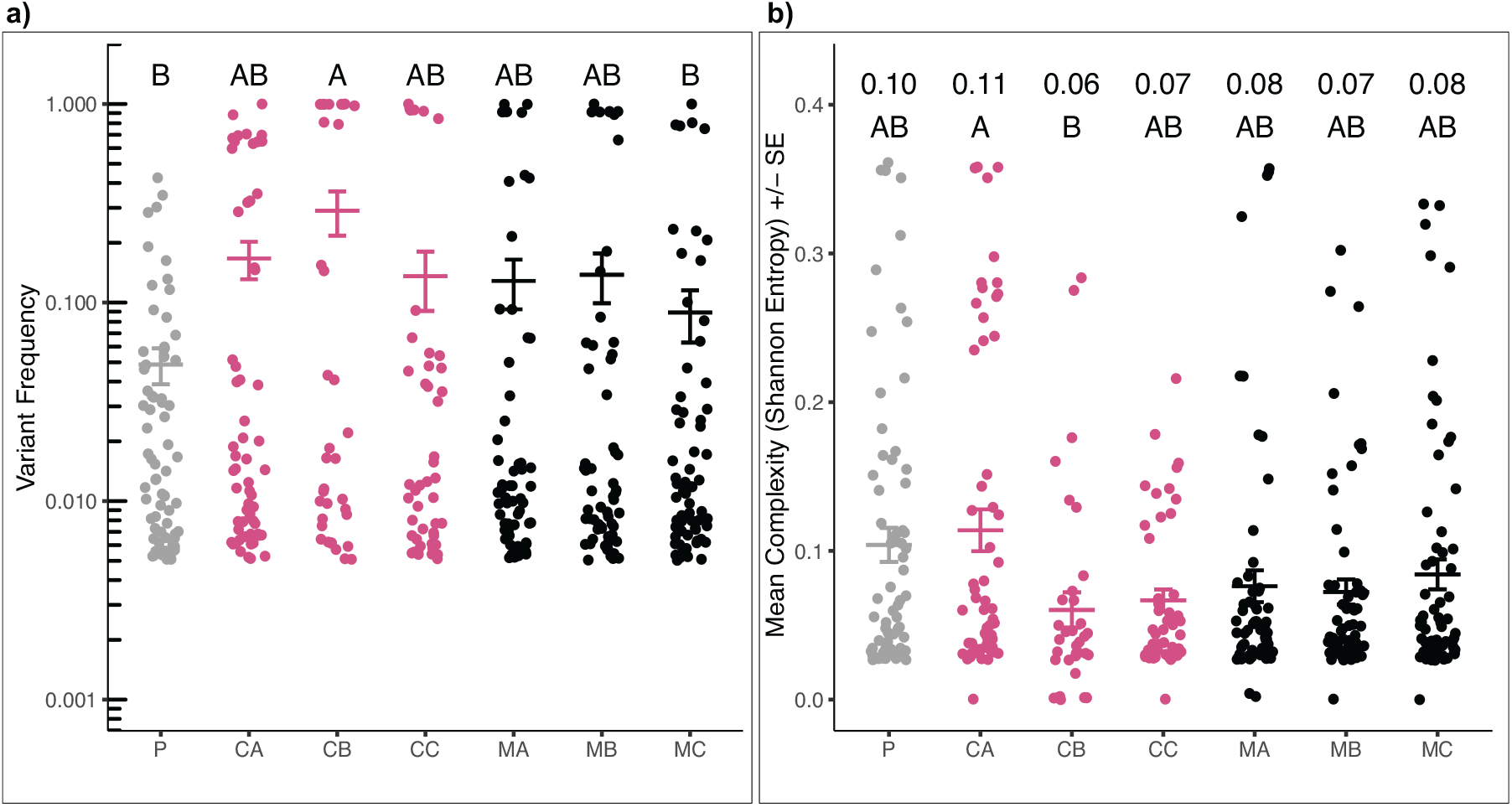
Ciprofloxacin-resistance does not alter frequency of DENV-4 variants or genetic complexity. a) The frequency of occurrence for DENV-4 variants from the parent (P; grey), ciprofloxacin-resistant (CA, CB, CC; pink), and media control (MA, MB, MC; black) populations was equivalent. b) Overall genetic complexity of ciprofloxacin-resistant DENV-4 (CA, CB, CC; pink) was not higher than the media control (MA, MB, MC; black) or parent (P; grey) populations. Mean Shannon Entropy values are indicated for each replicate. Differences were detected with ANOVA and Tukey pairwise comparisons (see text for full statistics).

**Figure 4.**
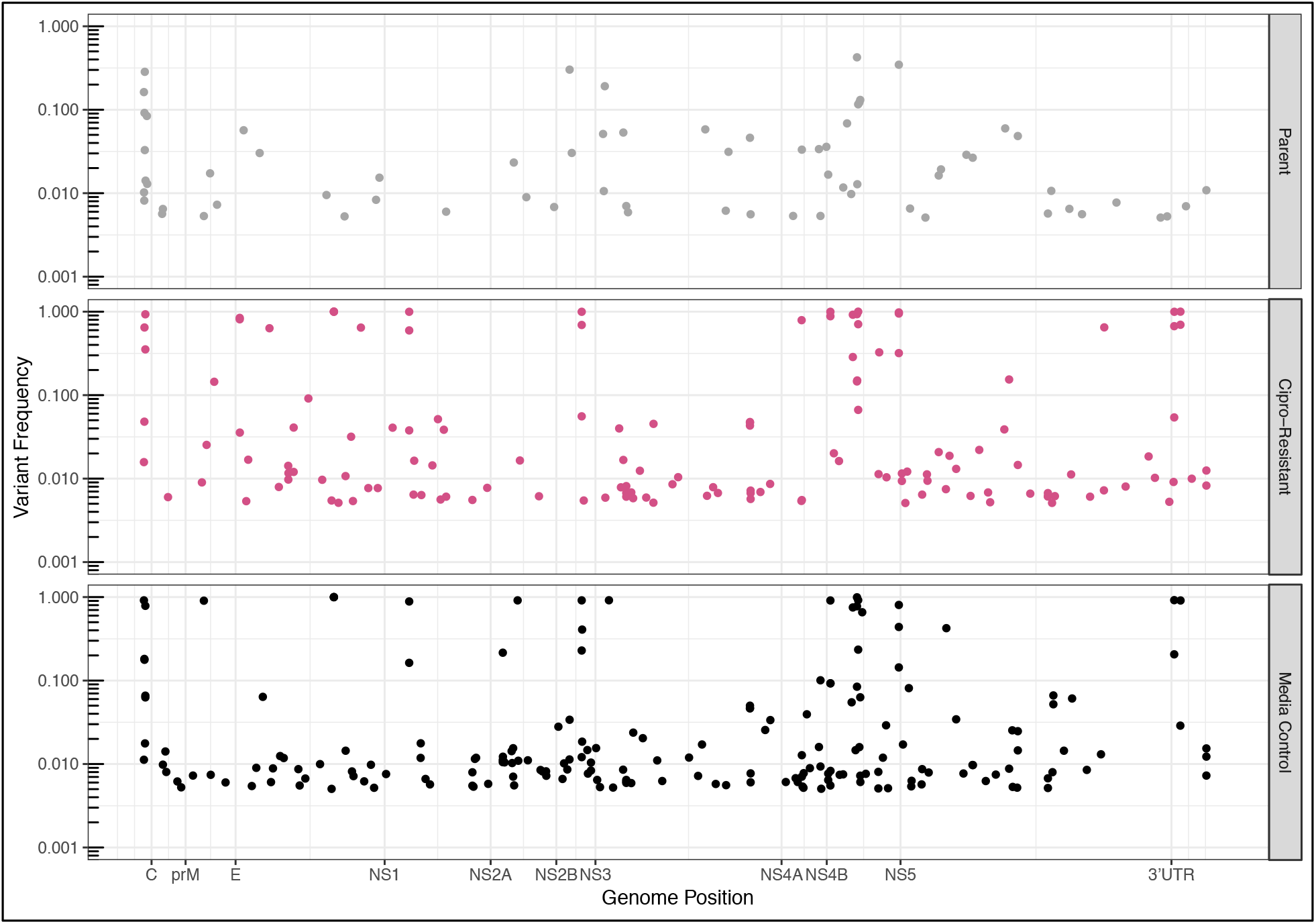
Variants in the ciprofloxacin-resistant and media-control populations are distributed across the DENV-4 genome. The frequency of occurrence for DENV-4 variants by genome position from the parent (grey), ciprofloxacin-resistant (pink), and media control (black). The start of each gene and untranslated region are indicated.

**Figure 5.**
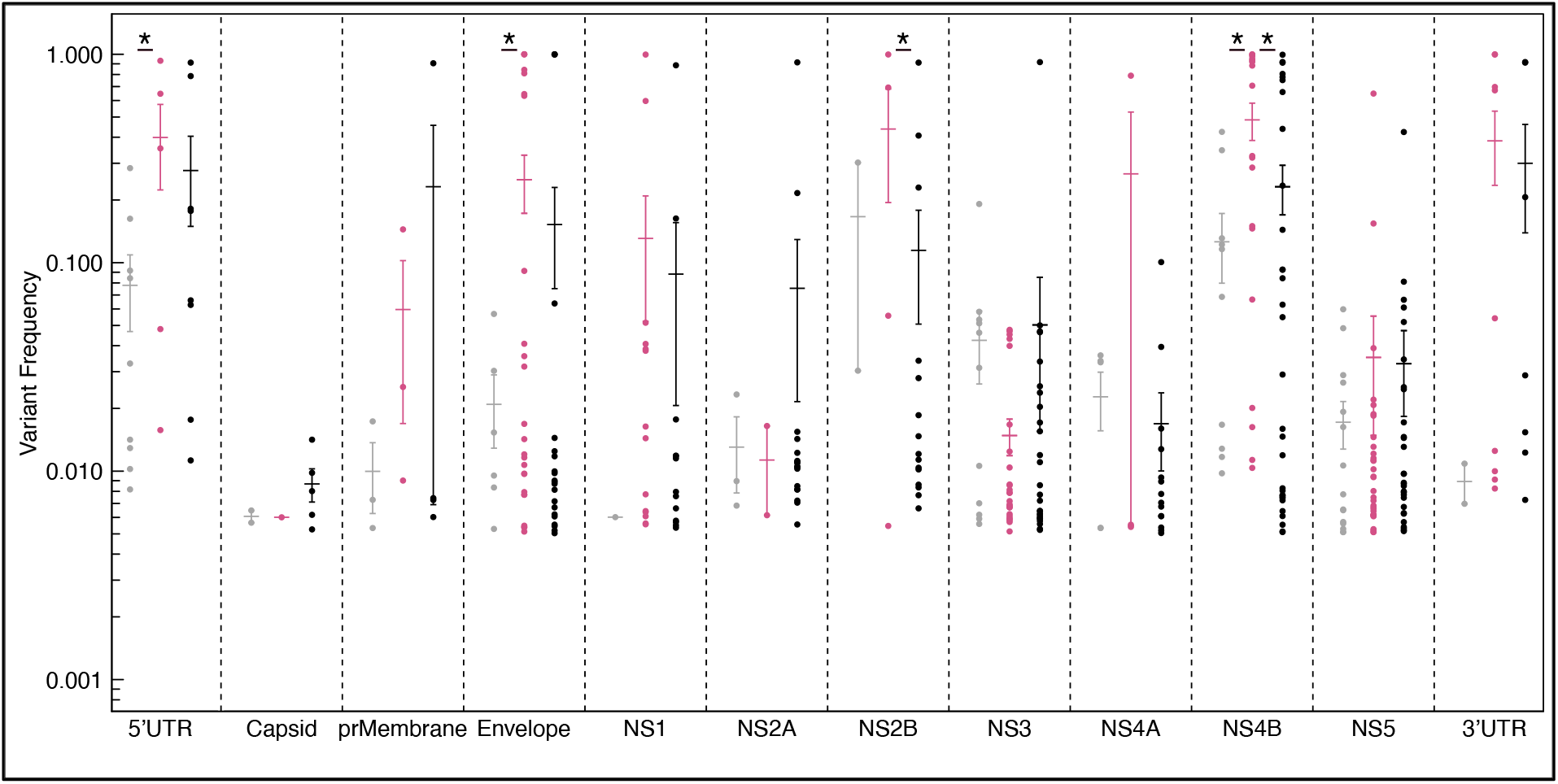
Ciprofloxacin-resistant populations experienced higher frequency of variants in the 5’ UTR, envelope, and NS4B. The frequency of occurrence for DENV-4 variants from the parent (grey), ciprofloxacin-resistant (pink), and media control (black) within each gene and UTR. Differences were detected with a two-way ANOVA (see text for statistics) and Tukey pairwise comparisons. Full pairwise statistics for differences within and between genes by treatment are in Table S2. * P < 0.05.

### Cross-resistance between ciprofloxacin and difloxacin does not extend to enoxacin

As shown in Figure 6a, viral replication of the fluoroquinolone-resistant and media control viruses in the presence and absence of the other fluoroquinolones revealed the efficacy of difloxacin and ciprofloxacin to suppress ciprofloxacin-resistant and difloxacin-resistant DENV-4, respectively, was diminished. Remarkably, enoxacin retained its ability to suppress viral replication of both ciprofloxacin- and difloxacin-resistant DENV-4. An interaction between viral lineage and treatment in the resistance test was detected using a two-way ANOVA (F (2,12) = 8.7, P = 0.005). Viral replication for both lineages decreased after treatment with enoxacin and difloxacin at their EC_50_ values, 7.6μM and 10.1μM, respectively (Tukey pairwise comparisons, P < 0.05). However, after difloxacin treatment, the ciprofloxacin-resistant DENV-4 replicated to the same level as the media control viruses treated with media. Viral replication after treatment with enoxacin remained 14x lower than the media control viruses treated with media, a highly significant difference (t = −4.3, df = 4. P = 0.01).

**Figure 6.**
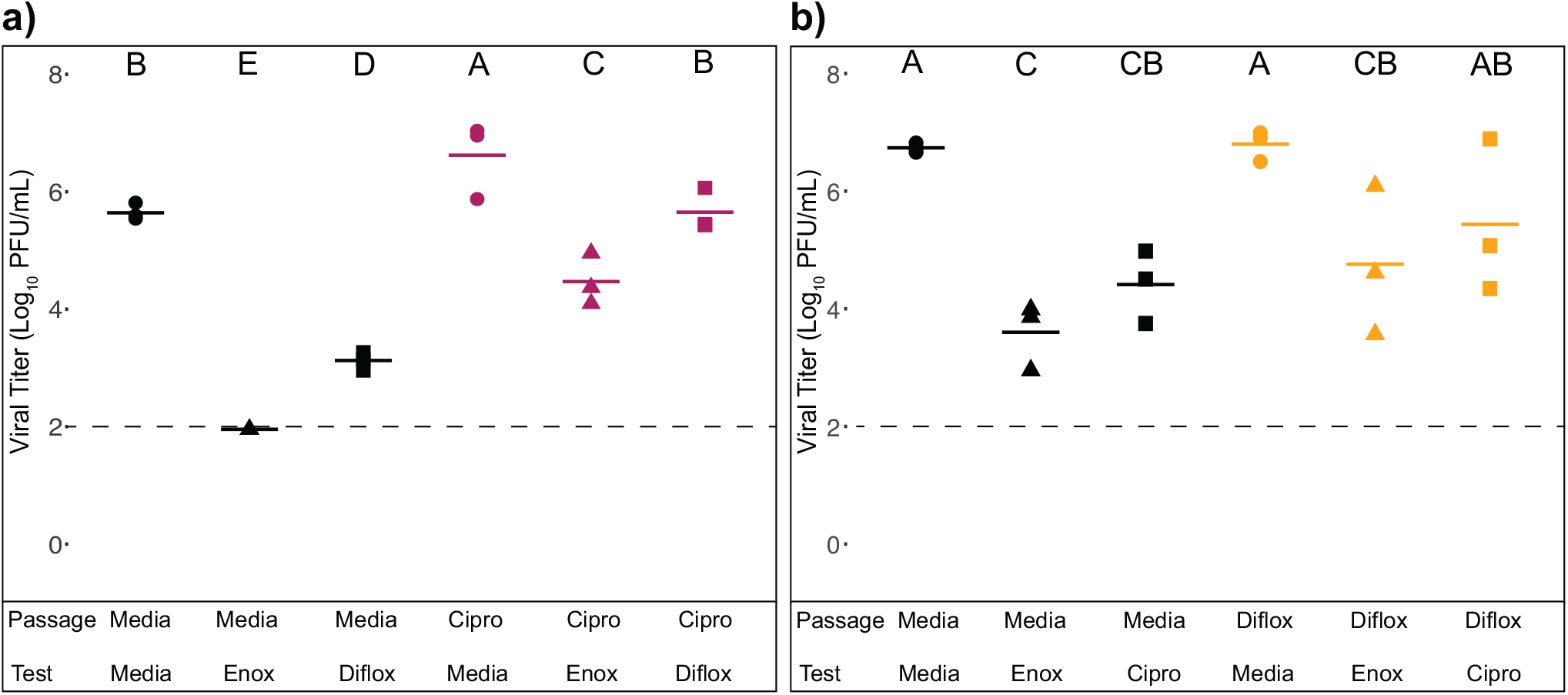
Ciprofloxacin-resistant DENV-4 was found to have cross-resistance to difloxacin, but not enoxacin (a) and difloxacin-resistant DENV-4 was found to have cross-resistance to ciprofloxacin, but not enoxacin (b). Black: media control passaged DENV-4, Pink: ciprofloxacin-resistant DENV-4, Yellow: difloxacin-resistant DENV-4, Circles: Tested in resistance test with media, Triangles: Tested in resistance test with enoxacin. Squares: Tested in resistance test with difloxacin (a) or ciprofloxacin (b). Differences were detected with two-way ANOVA with Tukey pairwise comparisons. Dashed line indicates the limit of detection.

A similar pattern was seen with difloxacin-resistant DENV-4 but interaction between viral lineage and treatment in the resistance test was not significant (two-way ANOVA F (2,12) = 0.8, P = 0.48; Figure 6b). The mean viral titer for the viruses passaged with media and tested against media were almost 100-fold higher than the mean viral titer for the difloxacin-resistant viruses tested with enoxacin, indicating that enoxacin still suppresses replication of DENV-4 that is resistant to difloxacin (Tukey pairwise comparison, P < 0.05). Yet, the mean viral titer of the difloxacin-resistant viruses tested with ciprofloxacin was not different from the media-passaged viruses treated with media demonstrating that DENV-4 resistance to difloxacin can result in cross-resistance to ciprofloxacin Tukey pairwise comparison, P > 0.05).

### In the absence of ciprofloxacin, ciprofloxacin-resistant DENV-4 are more fit than controls in two lines of human cells but not Vero or mosquito cells

Multicycle replication curves (Figure 7) reveal that, in the absence of ciprofloxacin, both control and ciprofloxacin-resistant DENV-4 gained fitness in the HEK-293 cells in which they were passaged relative to the parent virus, and that ciprofloxacin-resistant lineages reach equivalent peak titer to media-control lineages but declined in titer more slowly (Figure 7). Viral titers of the ciprofloxacin-resistant and media control DENV-4 were an average of 2.7 log and 2.5 log higher, respectively, than the parent from days 1 to 3 p.i. Repeated measures ANOVA detected a significant interaction between viral lineage and time (Figure 7; F (6, 24) = 6.1, P < 0.0006) and greater titer of the ciprofloxacin-resistant than media control DENV-4 on days 3, 4, and 6 p.i. were detected by a pairwise t-test *post hoc* test (P < 0.05).

**Figure 7.**
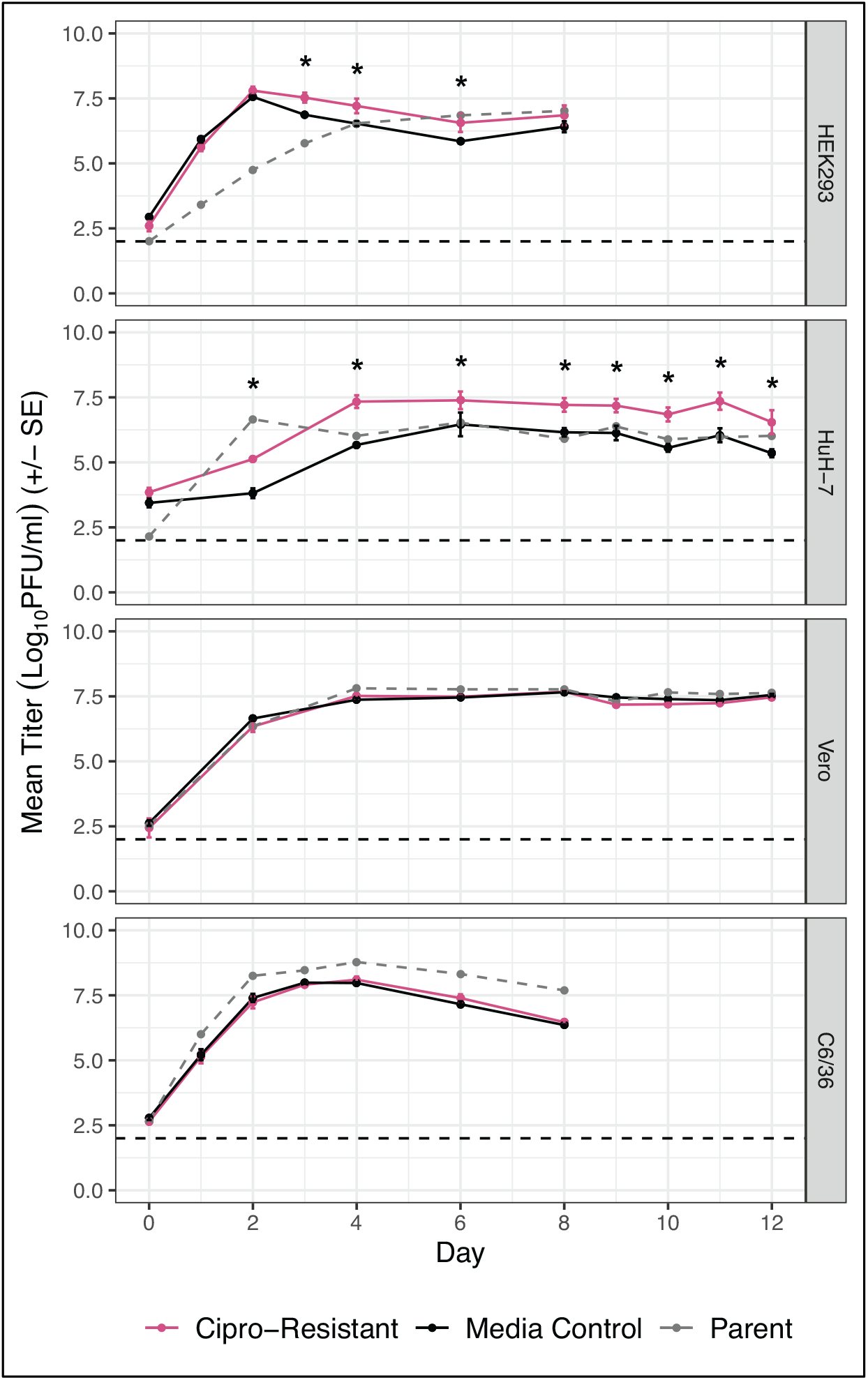
Replication kinetics of ciprofloxacin-resistant (pink solid line), media control (black solid line), and parent (grey dashed line) DENV-4 were different in HEK-293 and HuH-7 cells but not in Vero or C6/36 cells. Differences detected with repeated measures ANOVA and Tukey pairwise comparisons. Dotted line indicates the limit of detection. * P < 0.05.

In HuH-7 cells, replication of the ciprofloxacin-resistant DENV-4 reached a higher peak titer by day 2 post-infection than the media control and sustained that titer for the 12 days over which sampling was conducted (Figure 7). While the interaction between viral lineage and time was not significant (repeated measures ANOVA: F (8, 32) = 1.3, P = 0.26), pairwise comparisons detected greater viral titers for the ciprofloxacin-resistant viruses compared to the media controls on days 2 - 12 p.i. (P < 0.05).

In African green monkey kidney cells (Vero), replication of the parent virus, ciprofloxacin-resistant and media-control virus lineages were all remarkably similar. Again, the interaction between viral lineage and time was not significant (repeated measures ANOVA: F (8, 32) = 1.0, P = 0.43) but no pairwise differences were detected (Figure 7).

In contrast to mammalian cells, the parent DENV-4 replicated to higher levels in mosquito C6/36 cells compared to the passaged viruses. An interaction between viral lineage and time was not detected (repeated measures ANOVA: F (6, 28) = 0.6, P = 0.71) and no pairwise differences were identified as significant between the viral titers of the ciprofloxacin-resistant and media control viruses (Figure 7).

### Ciprofloxacin-resistant DENV-4 virions are more stable than controls

Over the course of 30 hours, both ciprofloxacin-resistant and media-passaged DENV-4 showed remarkably greater stability than the parent virus, and the ciprofloxacin-resistant viruses were also significantly more stable than media-passaged virus, although this difference only manifested at the last sampling point of the 30-hour test (Figure 8). A repeated measures ANOVA did not detect an overall interaction between percent change in viral titer and time (F (7,28) = 1.8, P = 0.13), but pairwise t-tests detected a significant difference at 30 hours between the ciprofloxacin-resistant viruses and media control, when mean viral titers for the ciprofloxacin-resistant viruses were 15% less than their initial values at time 0, whereas the media control viral titers were reduced by 52% (Figure 8).

**Figure 8.**
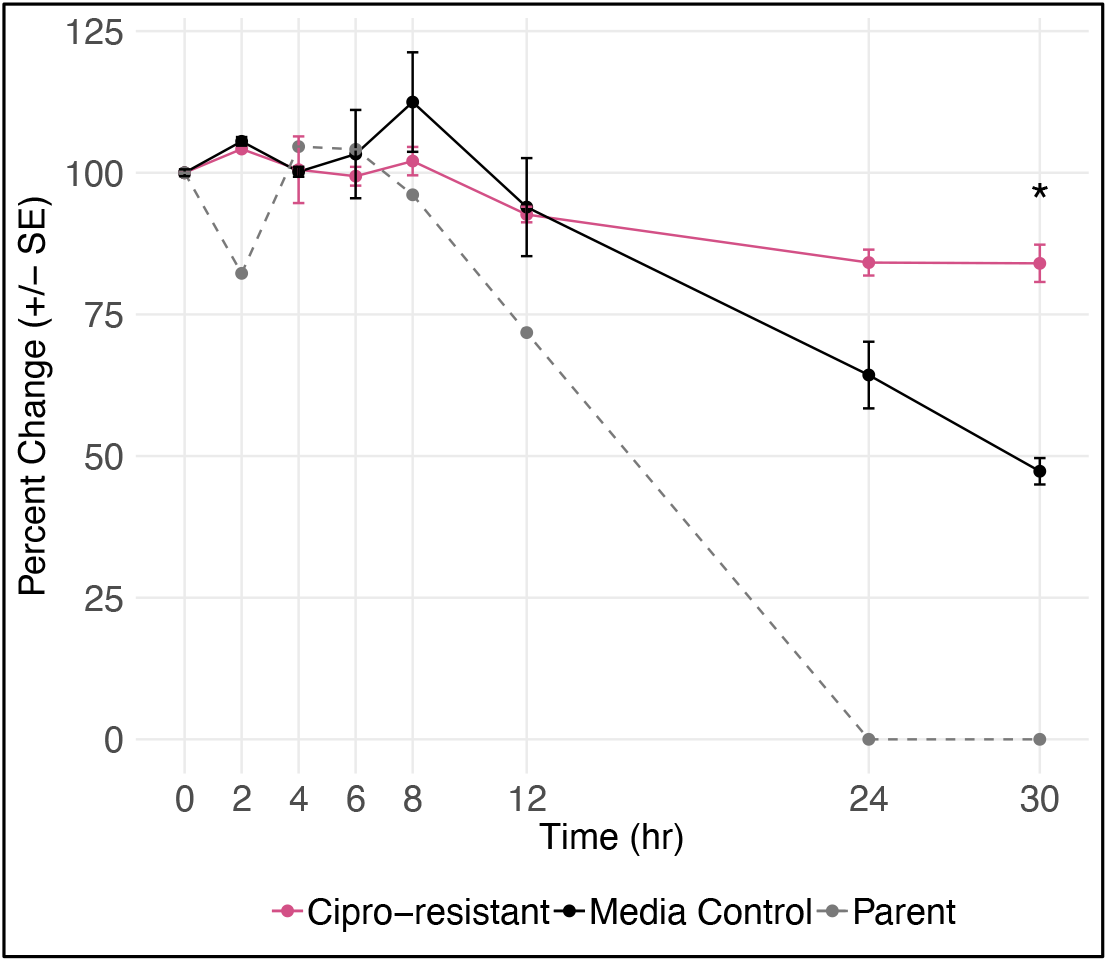
Virions of ciprofloxacin-resistant DENV-4 (pink solid line) were more stable than the media control (black solid line) and parent (grey dashed line) DENV-4 virions. Differences in the percent of initial viral titer were detected with repeated measures ANOVA and Tukey pairwise comparisons. * P < 0.05.

### Ciprofloxacin-resistant and media-control DENV-4 are unable to infect live mosquitoes

To evaluate the impacts of ciprofloxacin-resistance on potential for mosquito transmission, *Ae. aegypti* were fed on bloodmeals containing one of each of the three replicates of ciprofloxacin-resistant or media control or the single replicate of parent DENV-4 and infection was assessed 10 days post-feeding. Mortality ranged from 5.3% to 63.2% and was not different by virus; ANOVA F (2,6) = 1.3, P = 0.3); most mosquitoes that died simply failed to revive after being chilled for sorting. Of the survivors who fed on the parent virus 24.3% became infected with an average titer of 2.25 log10PFU/body, however, none of mosquitoes that fed on bloodmeals containing passaged virus (ciprofloxacin-resistant or media control) became infected (Table 3; χ^2^ = 22.0, df = 2, P < 0.0001). Of the infected mosquitoes that fed on bloodmeals spiked with parent virus, 45% of the heads were also infected with an average titer of 2.69 log10PFU/head (Table 3), while, as expected based on bodies, the heads of all the mosquitoes who fed on the ciprofloxacin-resistant or media control viruses were negative for DENV-4 infection.

**Table 3.**
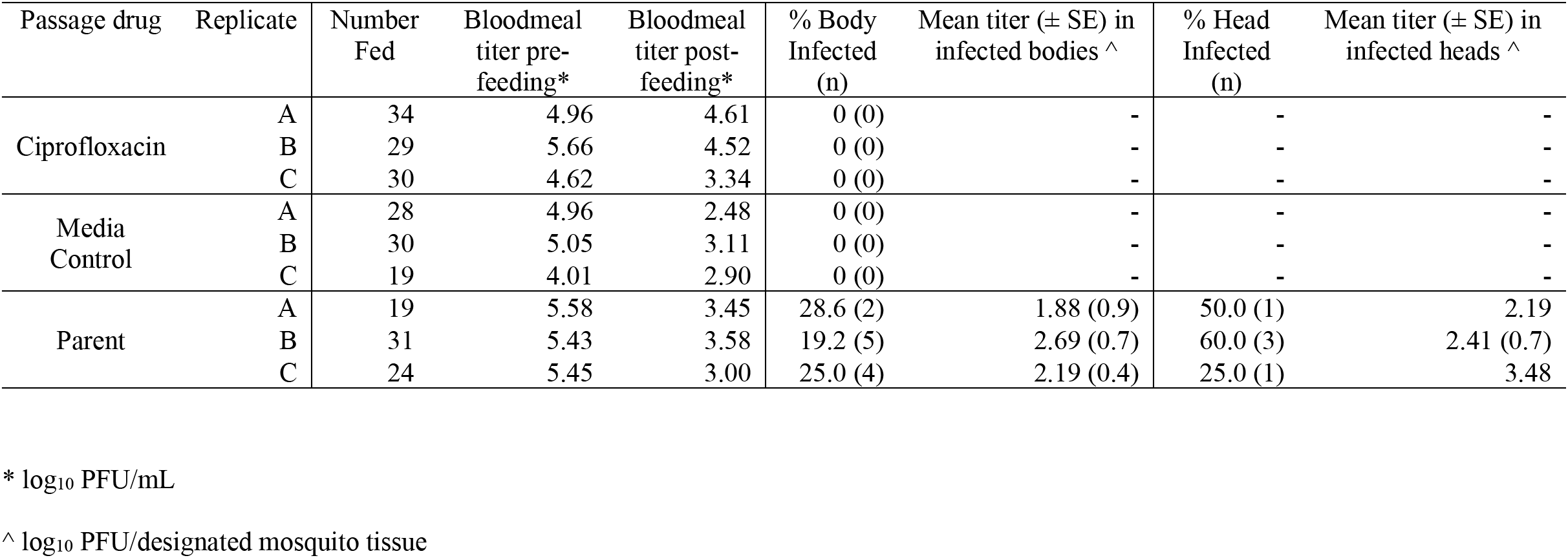
Infection of ciprofloxacin-resistant, media control, and parent DENV-4 in *Aedes aegypti* mosquitoes 10 days after feeding with an infectious artificial bloodmeal

| Passage drug | Replicate | Number Fed | Bloodmeal titer pre-feeding* | Bloodmeal titer post-feeding* | % Body Infected (n) | Mean titer (± SE) in infected bodies ^ | % Head Infected (n) | Mean titer (± SE) in infected heads ^ |
| --- | --- | --- | --- | --- | --- | --- | --- | --- |
| Ciprofloxacin | A | 34 | 4.96 | 4.61 | 0 (0) | - | - | - |
|  | B | 29 | 5.66 | 4.52 | 0 (0) | - | - | - |
|  | C | 30 | 4.62 | 3.34 | 0 (0) | - | - | - |
| Media Control | A | 28 | 4.96 | 2.48 | 0 (0) | - | - | - |
|  | B | 30 | 5.05 | 3.11 | 0 (0) | - | - | - |
|  | C | 19 | 4.01 | 2.90 | 0 (0) | - | - | - |
| Parent | A | 19 | 5.58 | 3.45 | 28.6 (2) | 1.88 (0.9) | 50.0 (1) | 2.19 |
|  | B | 31 | 5.43 | 3.58 | 19.2 (5) | 2.69 (0.7) | 60.0 (3) | 2.41 (0.7) |
|  | C | 24 | 5.45 | 3.00 | 25.0 (4) | 2.19 (0.4) | 25.0 (1) | 3.48 |

\* log<sub>10</sub> PFU/mL
^ log<sub>10</sub> PFU/designated mosquito tissue

## DISCUSSION

We recently demonstrated the efficacy of three fluoroquinolones, ciprofloxacin, difloxacin and enoxacin, against a panel of flaviviruses in cultured cells and A129 mice (5). In the current study, we investigated the barrier to evolution of resistance and cross-resistance to these drugs, their mechanisms-of-action, and the fitness consequences of resistance. DENV-4 was passaged in triplicate in the presence or absence of each drug until resistance was detected or to a predetermined endpoint of 10 passages.

We found that DENV-4 evolved resistance to ciprofloxacin within seven passages and difloxacin within ten passages but the virus failed to evolve resistance to enoxacin within ten passages, suggesting a higher barrier to resistance against enoxacin than the other two drugs. The variation in the rate of evolution of resistance could be due to stochasticity in evolution, particularly as our definition of resistance required that all three replicates within a treatment evolve resistance, but variation in resistance evolution could also reflect differences among the drugs in in mechanism-of-action. In our previous study of these drugs, time-of-addition assays revealed that enoxacin suppresses intermediate life cycle stages of ZIKV, such as translation of the polyprotein and replication of the genome while ciprofloxacin and difloxacin suppress ZIKV during both early and intermediate life cycles stages, suggesting that ciprofloxacin and difloxacin may share a mechanism-of-action that is different than that of enoxacin (5). Further supporting this contention, ciprofloxacin resistance conferred resistance to difloxacin and vice versa, but neither resistance to ciprofloxacin nor difloxacin conferred resistance to enoxacin. Differences in biochemical activities among the three drugs are documented (67); in particular, while enoxacin strongly enhances RNAi, ciprofloxacin and difloxacin have a moderate and marginal effect on RNAi, respectively (24). Cross-resistance does raise concerns about the long-term sustainability of ciprofloxacin and difloxacin as flaviviral inhibitors, but combination therapies with other antiviral compounds that are not similar in structure and do not share a mechanism-of-action may mitigate these concerns.

To gain further insight into the antiviral mechanism-of-action, we sequenced the whole genome of each of the three ciprofloxacin-resistant viruses, three media-control viruses, and the parent virus. We discovered two mutations, both in the envelope glycoprotein, that were unique to the ciprofloxacin-resistant viruses: V15L which arose in two replicates and E417A which arose in one replicate. E417A was also detected in one replicate of the difloxacin-resistant viruses, but neither mutation was detected in the enoxacin-passaged viruses. Ciprofloxacin-resistant viruses were sequenced by Illumina while difloxacin-resistant and enoxacin-passaged viruses were analyzed via Sanger sequencing, thus it is possible that the mutations detected in the ciprofloxacin-resistant lineages occurred in these two groups at frequencies below the level of detection. Also, resistance to difloxacin was not as complete as resistance to ciprofloxacin, as resistance was confirmed at the EC_50_ (10.1μM), but not at 2x EC_50_ (20.2μM). Envelope dimers form the herringbone protein coat of the virion (68). Upon acidification of the endosome, genome release is initiated when domain II folds out at the hinge point located between domain I and II, forming contact points with the cellular vesicle (69). V15L occurred in the envelope protein in domain I, a β-barrel that organizes two envelope proteins into a dimer (69), while E417A occurred in domain III. The conservative change of valine to leucine at amino acid position 15 is unlikely to cause a major change in protein structure, and indeed DENV serotypes 1 and 3 possess a leucine at this position. However, replacement of a negatively charged glutamic acid at amino acid with a hydrophobic alanine could potentially shift protein structure and function; all vector-borne flaviviruses analyzed possessed a negatively charged amino acid at this position and none of the 13 flaviviruses analyzed possessed an alanine. Additionally, a panel of 9 mutations were shared by resistant and control viruses, many of which have been previously shown to enhance replication in mammalian cells (40,70–73).

Contra our prediction about the likely role of helicase (NS3) in the action of fluoroquinolones, no consensus mutations were detected in NS3 in the ciprofloxacin-resistant viruses. This data indicates that ciprofloxacin does not interfere with DENV-4 helicase, as has been suggested for the inhibition of HCV by fluoroquinolones (26). Ciprofloxacin-resistant viruses also did not show increased variant frequency or genetic diversity; such an increased would be expected if ciprofloxacin suppressed viral replication by enhancing RNAi (74,75).

Our findings do suggest that that ciprofloxacin and difloxacin antagonize virus binding or entry. Fluoroquinolones are produced as a byproduct during chloroquine synthesis and share similarities in structure (Figure 1) (76), and the action of chloroquine may offer a clue about how ciprofloxacin and difloxacin could affect DENV entry. Chloroquine has been shown to suppress multiple viruses including DENV, ZIKV, chikungunya virus, and poliovirus (77–84) likely by blocking endosome acidification and thereby inhibiting virus entry (79,82,85,86). Interest in chloroquine as an antiviral recently spiked due to its potential utility against the novel coronavirus, SARS-CoV-2, that in 2020 has caused a massive and ongoing pandemic. Unpublished data suggests that like chloroquine, ciprofloxacin neutralizes the pH level within the lumen of the *trans*-Golgi network of bronchial epithelial cells (87). As antiviral assays are conducted to determine the efficacy of chloroquine to suppress coronaviruses, it would be prudent to evaluate the ability of fluoroquinolones to suppress coronaviruses as well.

Alternatively, mutations in envelope could increase the stability of the protein and thereby slow virion degradation. The envelope protein of ZIKV and the ZIKV virion as a whole are more stable and infectious at high temperatures than DENV (88); this thermostability of ZIKV depends on hydrophobic-hydrophilic interactions in the αβ helix of domain II of E (89). In this study, both ciprofloxacin-resistant and media control virions were dramatically more stable than the parent virion, and ciprofloxacin-resistant virions were slightly more stable than the media controls. Moreover, both ciprofloxacin-resistant and media control DENV-4 had higher stability than that reported for wild type ZIKV (90). Both ciprofloxacin-resistant and media control share a E327G mutation in domain III of E, which is the most likely cause of their enhanced stability. Añez et al. found that passage of DENV-4 in fetal rhesus lung (FRhL) cells resulted in the same E E327G mutation, and that DENV-4 carrying this mutation exhibited an increase in replication and infectivity in FRhL cells compared to its parent (73). To bind to a target cell the DENV E protein interacts with heparan sulfate (91) and the E327G mutation increases DENV-4 binding affinity for heparan sulfate (73). Molecular modeling further predicted that the E327G mutation increased the net positive charge on the surface of the virion and suggested that the mutation created a new heparan sulfate binding site (73). The increase in virion stability of the ciprofloxacin-resistant and media control DENV-4 is likely due to the increased positive charge on the surface of the virion. Importantly, when two rhesus macaques were infected with the DENV-4 passaged in FRhL cells, neither monkey had detectable levels of viremia or antibodies indicating that a glutamic acid at position 327 is favorable for monkey infection even if a glycine does increase infection, replication, and heparan sulfate binding in cell culture (73).

A third possibility is that V15L and/or E417A could increase overall fitness to overcome suppressive effects of ciprofloxacin. The fitness of ciprofloxacin-resistant, media-control and parent viruses in the absence of the drug was assessed in four different cell lines. Under these conditions, viral titers of ciprofloxacin-resistant viruses were greater than those of the media control viruses in HEK-293 and HuH-7 cells. This gain in fitness in the absence of ciprofloxacin contradicts our hypothesis that resistance evolved due to a specific drug evasion. Intriguingly, replication of the three virus lineages were indistinguishable in Vero cells, a cell line that lacks an interferon response (92,93). It is not clear from our data whether the gain in fitness seen in HEK-293 and HuH-7 cells, but not Vero cells is dependent on the lack of interferon and/or is an artifact of adaptation to human cells. Importantly, we also do not know how this gain in fitness will impact infection *in vivo*.

Further, fitness of the ciprofloxacin-resistant and media-control viruses were equivalent, but lower than that of the parent virus, in C6/36 cells, a mosquito cell line that lacks a functional RNAi response (94) and both lineages failed to infect live *Ae. aegypti* mosquitoes fed on artificial bloodmeals spiked with these viruses. The failure of the passaged viruses to infect mosquitoes is likely due to the suite of mutations in NS4B that they accrued; some of these mutations have previously been associated with mammalian adaptation and loss of mosquito infectivity (40,70–73). This finding is consistent with the trade-off hypothesis, which states that a gain in fitness in the mammalian host will negatively impact fitness in the mosquito vector (51–53). Our data underscore the complexity of viral fitness and how the patterns are likely influenced by a combination of drug resistance-associated and cell culture adaptation mutations.

Fluoroquinolones show some promise for repurposing as anti-flavivirals, but this study also suggests several lines of inquiry that should be pursued to further evaluate the consequences of evolution of fluoroquinolone resistance. While the barrier to resistance to enoxacin was relatively high, DENV-4 evolved resistance to ciprofloxacin and difloxacin within ten passages at or near the EC50 of each drug. Furthermore, resistance to either ciprofloxacin or difloxacin confer cross-resistance to the other drug. Unexpectedly, mutations associated with resistance occurred in the envelope gene of the virus; the phenotypic impact of these E mutations should be formally assessed using reverse genetics in a future study. These mutations were associated with substantial gains in virion stability and viral fitness, but the degree to which the increase in fitness is attributable to the envelope mutations associated with resistance and how the increase in fitness in human cells will play out *in vivo* remains unknown. FFuture experiments to interrogate evolution of resistance to these fluoroquinolones *in vivo* are needed to better assess their potential and perils as antiflaviviral therapies.

## AUTHOR CONTRIBUTIONS

**Stacey Scroggs**: Conceptualization, Methodology, Formal analysis, Investigation, Project administration, Validation, Data curation, Visualization, Writing - original draft, Writing - review & editing; **Jordan Gass**: Investigation, Writing - original draft, Writing - review & editing; **Ramesh Chinnasamy:**Resources, Writing - review & editing; **Steven Widen**: Investigation, Resources, Writing - review & editing; **Sasha R. Azar**: Investigation, Writing - review & editing; Shannan Rossi: Writing - review & editing; **Jeffrey Arterburn**: Funding acquisition, Resources, Supervision, Writing - review & editing; **Nikos Vasilakis**: Funding acquisition, Resources, Supervision, Writing - review & editing; **Kathryn Hanley**: Conceptualization, Methodology, Funding acquisition, Formal analysis, Validation, Supervision, Writing - original draft, Writing - review & editing

## ACKNOWLEDGMENTS

We would like to thank Brett Moehn for assistance with analysis.

## Supplemental

**Table S1.** Silent nucleotide mutations for coding region in the ciprofloxacin-resistant, media-control viruses from passage 7, as well as parent DENV-4

| Gene | Nucleotide change | Amino acid change | Present in ciprofloxacin-resistant DENV-4 (P7) |  |  | Present in media control DENV-4 (P7) |  |  | Parent DENV-4 |
| --- | --- | --- | --- | --- | --- | --- | --- | --- | --- |
|  |  |  | A | B | C | A | B | C |  |
| prM | T623A | Silent | - | - | - | X | - | - | - |
| E | T1277C | Silent | X | - | - | - | - | - | - |
| NS1 | A2669G | Silent | X | X | - | - | X | - | - |
| NS2A | G3749T | Silent | - | - | - | X | - | - | - |
| NS3 | A4658G | Silent | - | - | - | X | - | - | - |
| NS4A | C6578T | Silent | - | X | - | - | - | - | - |
| NS4B | T7088C | Silent | - | - | X | - | - | X | - |
| NS5 | T9593C | Silent | X | - | - | - | - | - | - |

-: Mutation not identified

**Table S2.**
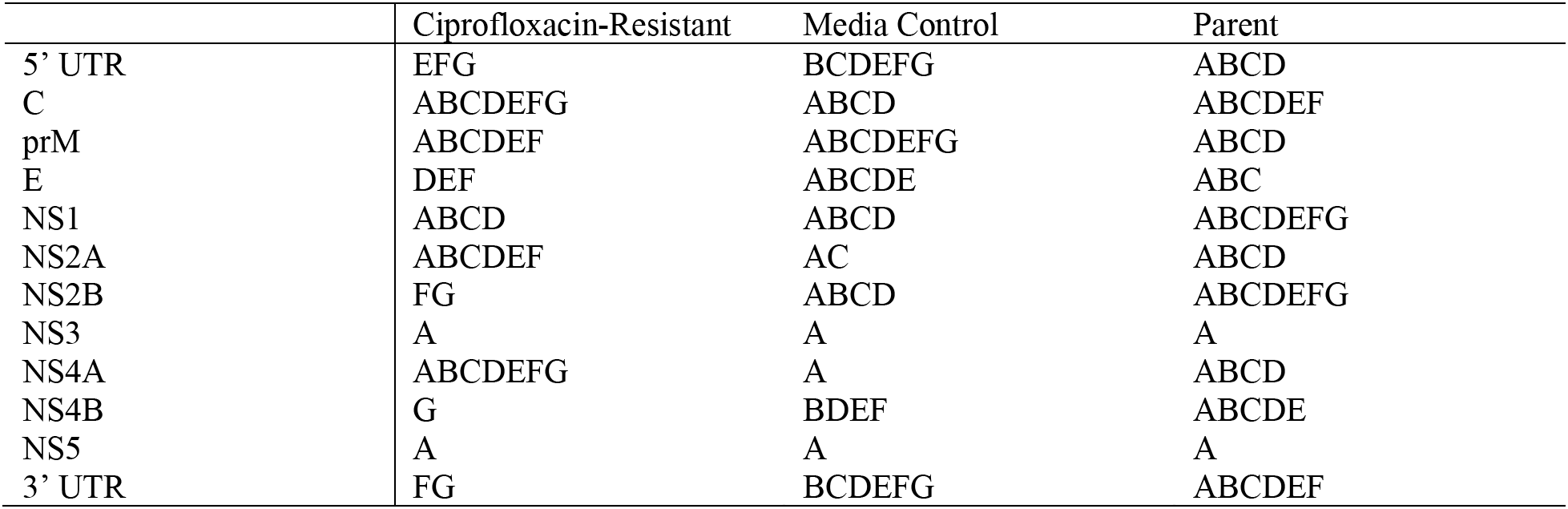
Full pairwise comparisons for differences between passage treatment condition and gene from Figure 5.

**Figure S1.**
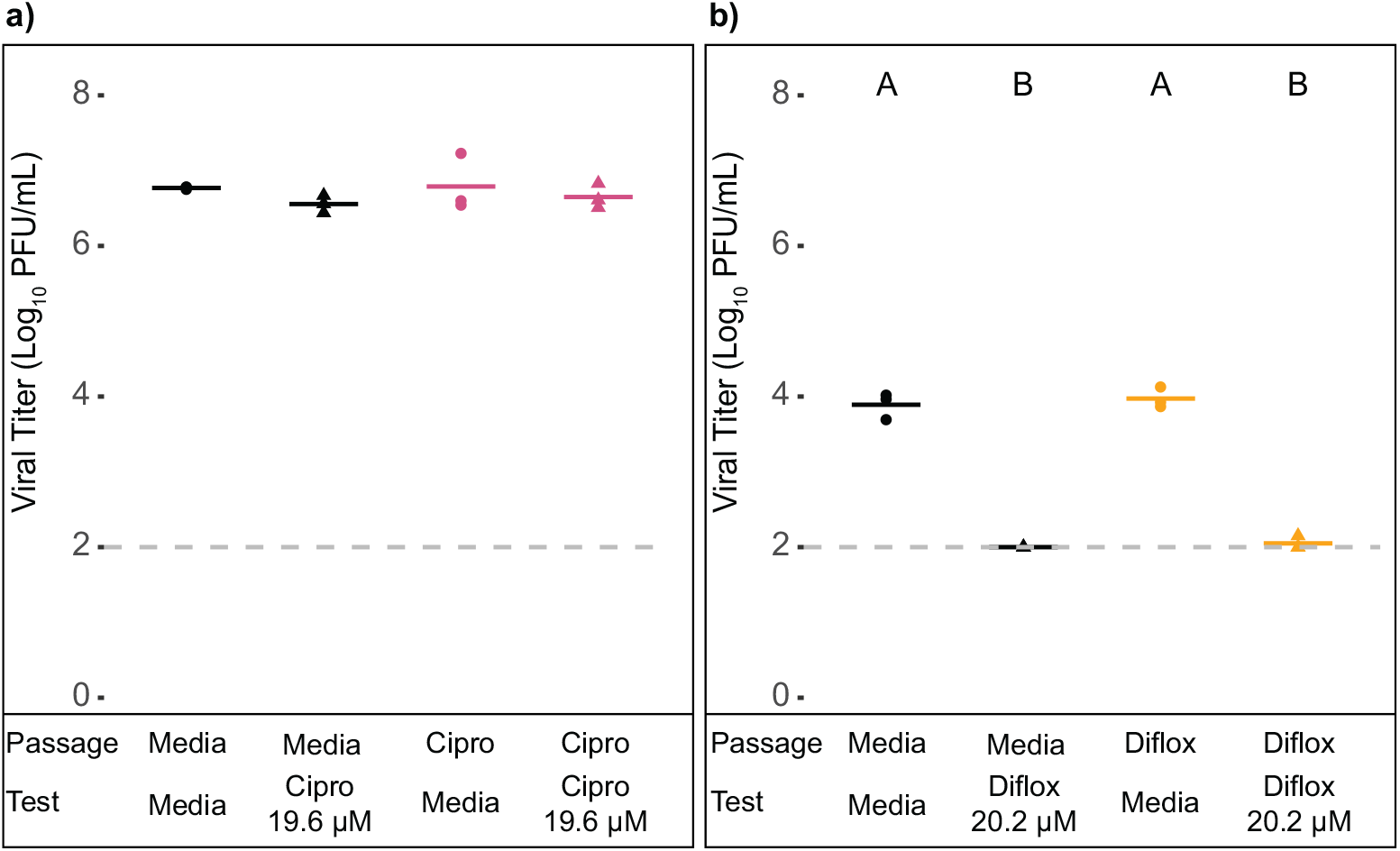
Viral replication of media-passaged, ciprofloxacin-passaged (a), and difloxacin-passaged (b) DENV-4 in the presence and absence of ciprofloxacin (19.6 μM) and difloxacin (20.2 μM). Black: media control passaged DENV-4, Pink: ciprofloxacin-passaged DENV-4, Yellow: difloxacin-passaged DENV-4, Circles: Tested in resistance test with media, Triangles: Tested in resistance test with 19.6μM (EC_50_) ciprofloxacin (a) and 20.2 μM (x2 EC_50_) difloxacin (b). Differences were detected with two-way ANOVA with Tukey pairwise comparisons. Solid lines indicate mean and dashed line indicates the limit of detection.

**Figure S2.**
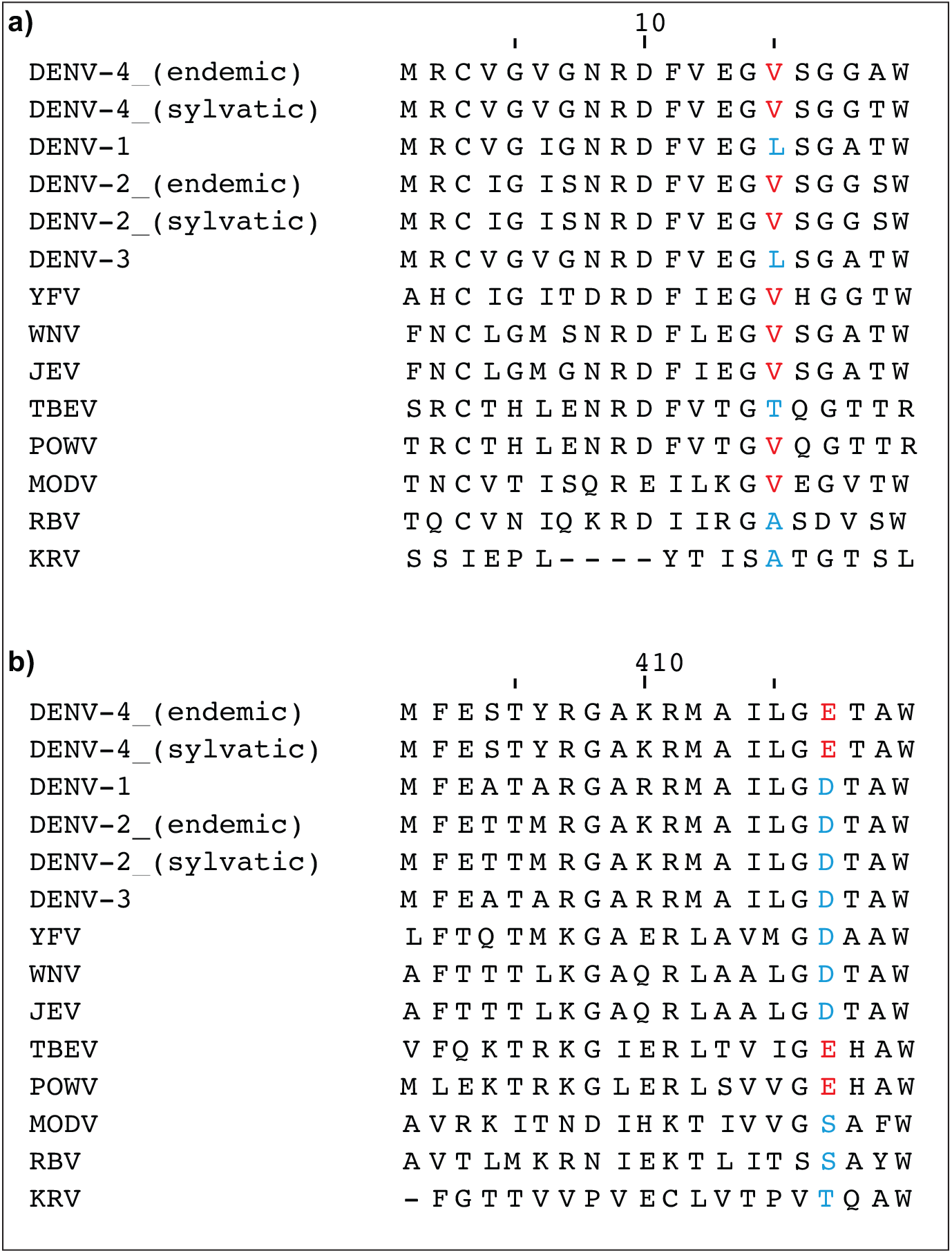
Sequence alignment of flavivirus envelope protein, a) amino acids 1-20 and b) amino acids 401-420. The amino acid sequences for the envelope gene were aligned for DENV-4 p4 (GenBank AY648301.1), DENV-4 p75 (GenBank EF457906.1), DENV-1 Western Pacific (GenBank AY145121.1), DENV-2 New Guinea C (GenBank AY243466.1), DENV-2 Dakar D75505 (GenBank EF457904.1), DENV-3 Sleman (GenBank AY648961.1), yellow fever virus RJ96 (GenBank MF423378.2), West Nile virus NY99 (GenBank HQ596519.1), Japanese encephalitis South Korea 2015 (GenBank MK541529.1), tick-borne encephalitis virus Sofjin (GenBank JN229223.1), Powassan virus PO375 (GenBank KU886216.1), Modoc virus (GenBank NC003635.1), Rio Bravo virus (GenBank JQ582840.1), and Kamiti River virus (GenBank NC005064.1) using MUSCLE v3.8 (95) and Jalview (96) v2.11. A blue amino acid at positions 15 and 417 indicate a change from the amino acid in DENV-4 at that position (red).

